# Structural variants contribute to phenotypic variation in maize

**DOI:** 10.1101/2024.06.14.599082

**Authors:** Nathan S. Catlin, Husain I. Agha, Adrian E. Platts, Manisha Munasinghe, Candice N. Hirsch, Emily B. Josephs

## Abstract

Comprehensively identifying the loci shaping trait variation has been challenging, in part because standard approaches often miss many types of genetic variants. Structural variants (SVs), especially transposable elements (TEs), are likely to affect phenotypic variation but we lack methods that can detect polymorphic structural variants and TEs using short-read sequencing data. Here, we used a whole genome alignment between two maize genotypes to identify polymorphic structural variants and then genotyped a large maize diversity panel for these variants using short-read sequencing data. After characterizing SV variation in the panel, we identified SV polymorphisms that are associated with life history traits and genotype-by-environment (GxE) interactions. While most of the SVs associated with traits contained TEs, only two of the SVs had boundaries that clearly matched TE breakpoints indicative of a TE insertion, while the other polymorphisms were likely caused by deletions. One of the SVs that appeared to be caused by a TE insertion had the most associations with gene expression compared to other trait-associated SVs. All of the SVs associated with traits were in linkage disequilibrium with nearby single nucleotide polymorphisms (SNPs), suggesting that the approach used here did not identify unique associations that would have been missed in a SNP association study. Overall, we have created a technique to genotype SV polymorphisms across a large diversity panel using support from genomic short-read sequencing alignments and connecting this presence/absence SV variation to diverse traits and GxE interactions.

## Introduction

A central question of evolutionary biology is how different types of mutations – single nucleotide polymorphisms (SNPs), insertion-deletion polymorphisms, copy number variants, translocations, and transposable element insertions – shape the phenotypic diversity observed in nature (Mitchell-Olds *et al*., 2007). Much recent effort has focused on characterizing structural variants (SVs): Tens of thousands of SVs have been identified in plant genomes (Darracq *et al*., 2018; Yang *et al*., 2019; Schatz, 2018; Alonge *et al*., 2020; Zhou *et al*., 2022; Qin *et al*., 2021; Hämälä *et al*., 2021) and specific SVs have been shown to affect important phenotypic traits in plants, including climate resilience in *Arabidopsis thaliana*, disease resistance and domestication traits in maize and rice, and frost tolerance in wheat (Beló *et al*., 2010; Cao *et al*., 2011; Sieber *et al*., 2016; Springer *et al*., 2009; Xu *et al*., 2012). In addition, maize SVs are predicted to be up to 18-fold enriched for alleles affecting phenotypes when compared to SNPs (Chia *et al*., 2012). These findings suggest that characterizing SV variation will be a crucial part of mapping genotypes to phenotypes.

A subset of SVs, transposable elements (TEs), are particularly interesting potential contributors to phenotypic variation (Lisch, 2013; Catlin and Josephs, 2022). TE content and polymorphism are shaped by a complex interplay of selection at the TE and organismal level (Charlesworth and Charlesworth, 1983; ^°^Agren and Wright, 2011) and there are many examples of TE variation affecting phenotypes (Hirsch and Springer, 2017; Lisch, 2013). For example, a TE insertion in the regulatory region of the *teosinte branched1* (*tb1*) gene in maize enhances gene expression causing the upright branching architecture in maize compared to its progenitor, teosinte (Studer *et al*., 2011). TE insertions also affect flesh color in grapes and fruit color and shape in tomato (Fray and Grierson, 1993; Kobayashi *et al*., 2004; Van der Knaap *et al*., 2004; Shimazaki *et al*., 2011; Domínguez *et al*., 2020). These phenotypic effects may result from changes in gene expression: TE activation can disrupt or promote gene expression (Hirsch and Springer, 2017; Fueyo *et al*., 2022), and the industrial melanism phenotype in British peppered moths, *Biston betularia*, results from TE-induced overexpression of a gene responsible for pigment production (Hof *et al*., 2016). TEs often activate (i.e. express and/or mobilize) in response to stress in many eukaryotes, including maize (Makarevitch *et al*., 2015; Liang *et al*., 2021), *Arabidopsis* (Wang *et al*., 2022; Sun *et al*., 2020), and *Drosophila melanogaster* (de Oliveira *et al*., 2021; Milyaeva *et al*., 2023), suggesting that they may contribute to trait variation in stressful environments. However, we lack systematic studies of how TEs in general affect phenotypic variation or how TEs may contribute to genotype-by-environment interactions outside of the context of stress.

Characterizing genomic variation for SVs and TEs has been challenging, especially in highly repetitive plant genomes where it is often difficult to uniquely align short-reads to the reference genome. Recent studies have shown that attempts to assemble SVs solely with short-read sequencing data can greatly underestimate the total number of SVs present in a population (Huddleston *et al*., 2017; Audano *et al*., 2019; Cameron *et al*., 2019; Ebert *et al*., 2021). Some estimates for the accuracy of SV discovery with short-read sequencing are as low as 11% in humans due to the inability of short-reads to align within highly repetitive regions, span large insertions, or concordantly align across SV boundaries (Lucas Lledó and Ćaceres, 2013). However, recent efforts using short-read sequencing from a population of grapevine cultivars have been used to genotype SVs by ascertaining SV polymorphisms between two reference genomes and calling these SVs within the population (Zhou *et al*., 2019).

The increasing availability of long-read sequencing has opened up an opportunity to identify SVs that would have been missed using short-read data. For example, long reads have been used to identify structural variants associated with traits in a set of 100 tomato accessions that were long-read sequenced (Alonge *et al*., 2020). In other systems without enough long-read sequenced genotypes to directly look for associations between structural variants and phenotype, researchers have started with SVs detected in a smaller subset of individuals with reference assemblies and then genotyped a larger mapping panel of individuals with short-read sequencing data. Researchers have used pan-genome graph methods to identify SVs in a smaller number of reference sequences and then genotype in a larger sample of short-read sequenced genotypes in *Arabidopsis thaliana* (Kang *et al*., 2023), soybean (Liu *et al*., 2020), rice (Qin *et al*., 2021), and tomato (Zhou *et al*., 2022). These studies have confirmed that SVs are important for trait heritability (Zhou *et al*., 2022). However, graph genome approaches are challenging for plants with large genomes and have not yet been widely adopted. For example, a haplotype graph has been generated for 27 maize inbred lines, but not for a wider diversity panel (Franco *et al*., 2020). Additionally, work using short-read alignments and pan-genome approaches have identified SVs in maize and found that SVs contributed to trait heritability (Gui *et al*., 2022). Approximately 60% of these SVs were “related” to TEs but no clear links between SV polymorphisms and TE insertions were made (Gui *et al*., 2022). Plants with large genomes are not only important for a number of practical reasons, but they also may have different genetic architectures underlying trait variation that evolve differently (Mei *et al*., 2018), so understanding how SVs and TEs contribute to trait variation in large-genomed plants is key for comprehensively understanding the importance of these variants in general.

To address the gap in understanding how SVs and TEs contribute to trait variation in a species with a large genome, we identified SVs found from the alignment of two reference assemblies using short-reads that overlap the SV junctions. This type of approach has been used previously in in a few other systems (Wang *et al*., 2020; Zhou *et al*., 2019). Here, we investigated the relationship between SV variation and phenotype in a diverse set of maize inbred lines in the Buckler-Goodman association panel (Flint-Garcia *et al*., 2005). After identifying SVs that differ between two accessions, B73 and Oh43, we genotyped 277 maize lines present in a larger mapping panel for the SV alleles. We detected SV polymorphisms that varied across the panel and linked these polymorphisms to phenotypic variation, GxE, and gene expression.

## Materials and methods

### Structural variant identification

An “ascertainment set” of SVs that differ between B73 and Oh43 were identified by Munasinghe *et al*. (2023). These genotypes were chosen to call SV presence/absence because they are both in the Buckler-Goodman association panel but come from different germplasm pools (Gage *et al*., 2019). Ascertainment set SVs were filtered to only contain those that had 300 bps of colinear sequence determined by AnchorWave (Song *et al*., 2022) in the immediate upstream and downstream regions flanking SV junctions. The apparent insertion and 300 bp flanking region on either side were extracted to create a FASTA file containing “SV-present” alleles. The corresponding site in the other genome where the SV was absent and 300 bp flanking sequences were also extracted and combined in the final FASTA file to serve as the “SV-absent” allele sequence. Ultimately, this FASTA file was used as a set of pseudoreference alleles to call SV polymorphism in individuals with only short-read sequence data (Figure S1).

### SV presence/absence genotyping

To call presence or absence for each SV, we collected genomic short-read data for 277 inbred maize genotypes from the Buckler-Goodman association panel sequenced for the third generation maize haplotype map (HapMap3) and aligned to the generated FASTA files with SV present and absent alleles (Flint-Garcia *et al*., 2005; Bukowski *et al*., 2018). Illumina adapters and low quality sequences were removed using Trimmomatic v0.39 (Bolger *et al*., 2014). PCR duplicate reads were also filtered out using the -r option within the *markdup* function in SAMtools v1.15.1 (Danecek *et al*., 2021). Surviving paired-end reads were merged into a master FASTQ file for each genotype and aligned to pseudoreference alleles using HISAT2 (Sirén *et al*., 2014). The aligned dataset was filtered to only contain concordant, uniquely mapping reads. We used read-depth for each upstream and downstream SV boundary to support the presence or absence of SVs (Figure 1). Read coverage at each SV boundary was calculated using the *coverage* function within bedtools v2.30.0 (Quinlan and Hall, 2010).

**Figure 1.**
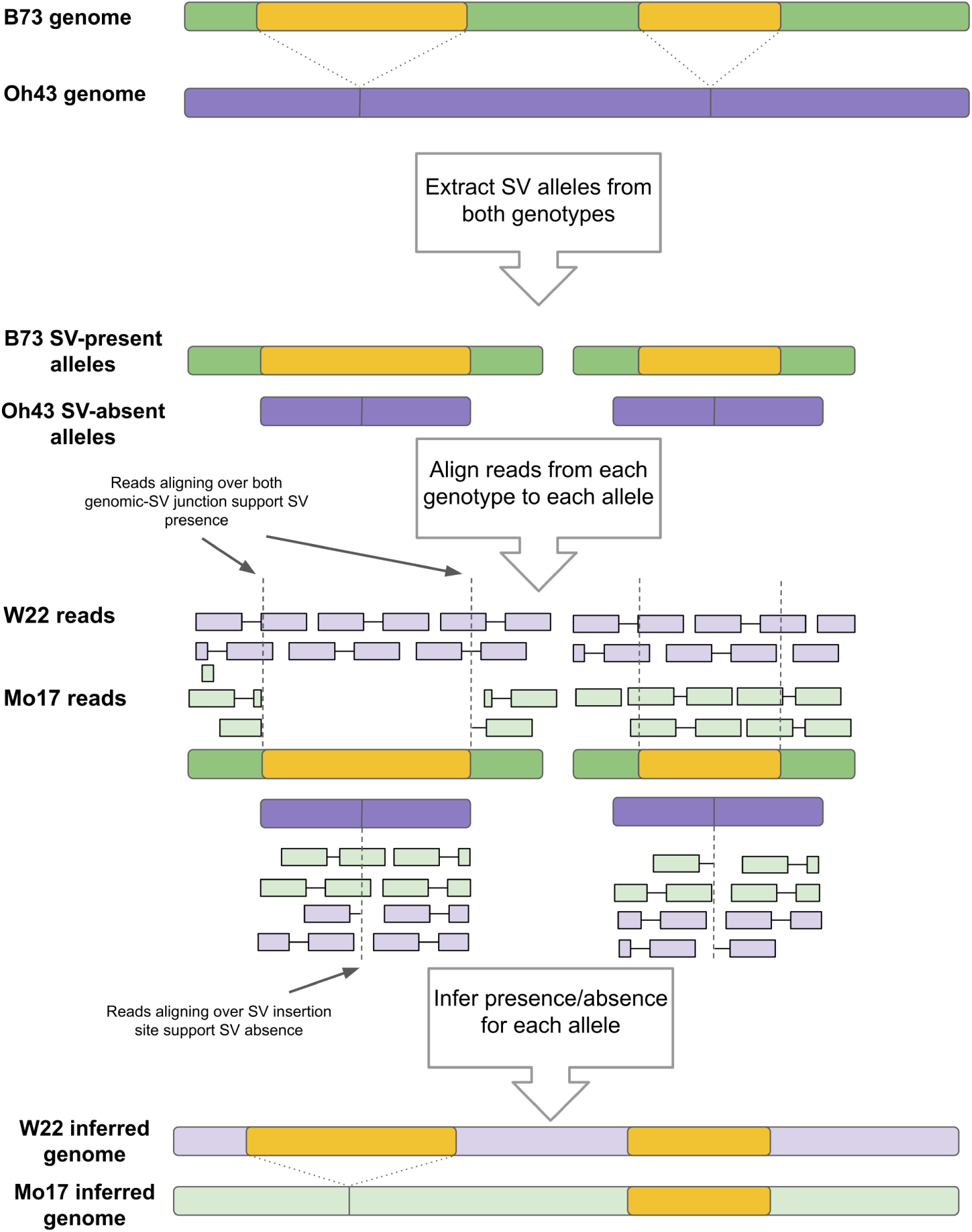
Method to call SV presence/absence with short read genomic data. – Using B73 and Oh43 as our ascertainment set, we first find polymorphic SVs between these two genotypes. To significantly improve read-mapping runtimes, we extract SVs and adjacent genomic sequences where SVs are present, while extracting only adjacent genomic regions at the polymorphic site where the SV is absent in the opposite genotype — termed pseudoreference SV alleles. Next, reads from a genotype of interest are mapped to these generated sequences. SVs can then be inferred present or absent based on their alignment to either allele.

First, we filtered out SVs where we were unable to use short-read data from B73 and Oh43 to correctly identify SV genotypes. In these cases, short-read data mapped better to the opposite genotype’s alleles than their own alleles. For an SV within our ascertainment set to be retained for downstream genotyping in the Buckler-Goodman association panel, we required that: (1) upstream and downstream SV junctions had the same or higher read coverage from the genotype with the SV than the other genotype and (2) no reads from the SV-present genotype spanned the insertion site for the genotype without the SV (Figure S2).

For the rest of the genotypes in the Buckler-Goodman association panel, SV-presence was supported in the query genotype if there was at least one read spanning the upstream or downstream SV junction and there was no read coverage at the SV polymorphic site for the alternative SV-absent allele. An SV-absent allele is supported if at least one read spans across the SV polymorphic site but no reads map to either SV junction of the corresponding SV-present allele. SVs are ambiguous if reads from the query genotype map to both the SV-present allele junctions and the SV-absent insertion site.

### Calculating linkage disequilibrium between SNPs and SVs

SNPs in variant call format (VCF) were collected from the third generation maize haplotype map version 3.2.1 and coordinates were converted to the B73 NAM reference positions (version 5) using liftOverVCF in Picard tools (Pic, 2019; Qiu *et al*., 2021a). Chain files for the genome builds B73 version 3 (APGv3) to B73 version 4 (B73 RefGen v4) and B73 version 4 to B73 version 5 (Zm-B73-REFERENCE-NAM-5.0) can be found in gramene.org and maizegdb.org, respectively (Tello-Ruiz *et al*., 2022; Woodhouse *et al*., 2021). We removed SNPs with *>* 10% missing data, a minor allele frequency (MAF) *<* 10%, and those within SV regions, resulting in 16,435,136 SNPs in the final filtered dataset. Additionally, we appended polymorphic SV calls for each genotype in the HapMap3 dataset to the final VCF file. Because SV-present alleles were characterized for both B73 and Oh43, we used the start of the SV coordinate for SV-present alleles within B73 and the B73 insertion site for SVs present in Oh43 as the coordinate for LD analysis. Following methods from Qiu *et al*. (2021a), we calculated LD between SNPs and nearby polymorphic SVs being sure to exclude SNPs inside of SVs, using PLINK v1.9 (Chang *et al*., 2015), www.cog-genomics.org/plink/1.9/ with the following parameters: --make-founders, --r2 gz dprime with-freqs, --ld-window-r2 0, --ld-window 1000000, --ld-window-kb 1000, and --allow-extra-chr.

### Association mapping

Polymorphic SVs across all query genotypes were converted to BIMBAM mean genotype format (Servin and Stephens, 2007). SV-present alleles that were characterized as ambiguous were denoted as NA. We performed a genome wide association (GWA) of SV presence/absence variants (PAVs) using phenotypes from Peiffer *et al*. (2014) and Bukowski *et al*. (2018), with a linear mixed model (LMM) in GEMMA v0.98.03 (Zhou and Stephens, 2012). The traits tested were collected from Peiffer *et al*. (2014) and are best linear unbiased predictions of the following: growing degree days to silking, growing degree days to anthesis, anthesis-silking interval measured in growing degree days, days to silking, days to anthesis, anthesis-silking interval measured in days, plant height, ear height, difference of plant height and ear height, ratio of ear height and plant height, and ratio of plant height and days to anthesis. To account for missing genotypic data for each SV, we required at least 90% of the genotypes to have presence/absence calls for relatedness matrix calculations and subsequent associations. All plots with genomic locations are shown with B73 coordinates, and Oh43 SV-present alleles were converted to B73 coordinates for display. To account for multiple-testing, we calculated a false discovery rate (FDR) adjusted significance threshold (Benjamini and Hochberg, 1995) to maintain an overall *α* = 5% significance. Filtered SNPs from the HapMap3 dataset were also subjected to GWA using the same methods as our polymorphic SV dataset.

In addition to the association analyses for main effects, we examined these data for genotype-by-environment interaction (GxE). For the 11 traits above, we used simple linear regression following the form of Finlay-Wilkinson (FW) regression (Finlay and Wilkinson, 1963) to record the slope (i.e. reaction norm) and mean squared error (MSE) for each genotype using the linear model (lm) function in R;

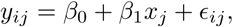

where *β*_0_ and *β*_1_ are the intercept and slope estimates for the *i^th^* line, respectively, *x_j_* is the average performance of all lines in the *j^th^* environment, and *ɛ_ij_* is a random error term. We removed any lines which were not represented in at least 6 environments on a per trait basis to reduce the error in our estimates. This filtering resulted in a different number of individuals and markers used in each FW model (ranging from 245 to 274 individuals per trait). We then performed GWA of SV PAVs using slope and MSE estimates for each trait as quantitative phenotypes in GEMMA as before.

### Gene expression

We used previously collected gene expression data for ∼37,000 maize genes (Kremling *et al*., 2018) to test for differential gene expression between SV genotypes at the loci identified in the association mapping analyses. We compared expression between SV genotypes for three tissue types: the tip of germinating shoots, the base of the third leaf and the tip of the third leaf. Library sizes were normalized using DESeq2 (Love *et al*., 2014) and we filtered the gene set to contain only genes with expression in 70% of individuals above 10 reads per median library size (approx 0.5 counts per million) using the edgeR package in R (Robinson *et al*., 2010), resulting in an average of 12,703 genes per SV identified in the GWAS. Finally, we used edgeR to test for differential expression by first building generalized linear models to model expression between genotypes and then testing for significance using the F-test. P-values were adjusted using FDR to maintain an overall significance threshold of *α* = 5%.

## Results

### Polymorphic SVs in the diversity panel

We genotyped SV polymorphisms for 277 maize genotypes at SVs segregating between B73 and Oh43 by aligning short reads from the genotypes to each SV allele and counting reads spanning genomic-SV junctions and SV polymorphic sites. Out of 98,422 polymorphic SVs between B73 and Oh43, we filtered out SVs where short reads from B73 and Oh43 did not clearly align to the correct allele. After this filtering step, we were able to determine the genotype of 64,956 SVs in the Buckler-Goodman association panel (Figure S2). The largest proportion of these SVs were those classified as “TE = SV” (21,103, 32.5%), followed by “multi TE SVs” (18,326, 28.2%), “incomplete TE SVs” (10,928, 16.8%), “no TE SVs” (8,842, 13.6%), and “TE within SVs” (5,757, 8.9%) (Figures S3, S4). The proportions of SVs for each category are consistent with those prior to filtering. For more information about how SVs are classified into TE groupings, see Munasinghe *et al*. (2023).

For subsequent analyses, we filtered the SV dataset to only include variants with a minor allele frequency (MAF) ≥ 10% and presence/absence calls for at least 90% of genotypes, resulting in the retention 3,087 SV alleles (4.75% of dataset) (Figure S5). Filtering on missing data and MAF removed many SVs because many individuals in the dataset have low realized sequencing coverage when mapped to the B73 reference assembly. There is a median coverage of 2.68, ranging from 0.031 in the A554 genotype to 19.47 in B57. Read depth per individual was negatively correlated with percent missing SV data per individual (*p* = 2.4 × 10^−5^) (Figures S6, S7), suggesting that missing data for SVs results from not having enough reads covering the junction sites. This pattern suggests that this method needs a minimum of average read depth of 5 to successfully genotype SVs at most sites, although this number will likely vary by species.

We investigated the frequency spectrum of SV polymorphisms in the Buckler-Goodman association panel by calculating the frequency of the allele with a putative insertion (or lacking a putative deletion). Since these SVs were initially identified as being polymorphic between two individuals, it was not surprising to see that many of the SVs were at moderate frequency in the population (Figures 2, S3). For most SVs, the SV-present allele was more common than the SV-absent allele. This pattern is consistent with the polymorphism being caused by a deletion and the longer ‘insertion’ allele being the ancestral type, and so present at higher allele frequencies in the population. The frequency spectrum was relatively consistent across SV types (Munasinghe *et al*., 2023).

**Figure 2.**
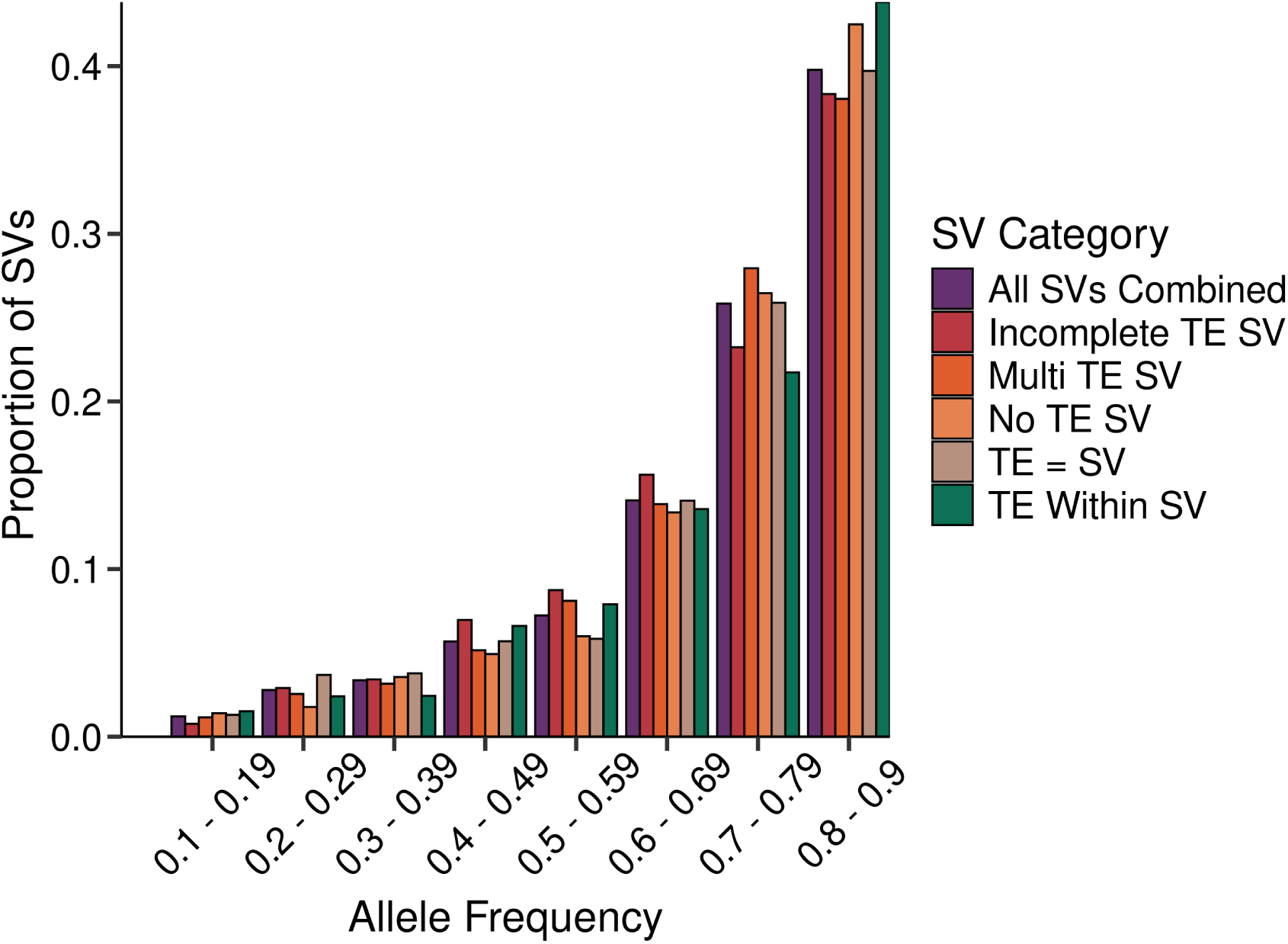
Site-frequency Spectrum of SVs. – SVs were filtered to only contain those with a minor allele frequency ≥ 10% and ≤ 10% missing data (n = 3,087). The SFS is unfolded and displays the frequency of the allele with the putative insertion (or that is lacking a deletion).

### SV genotypes are associated with phenotypic traits

In a genome-wide association analysis, SV presence/absence was significantly associated (FDR *<* 0.05) with four out of the eleven traits tested: growing degree days to anthesis, days to silking, days to anthesis, and ear height (Figures 3, S8). All four SV associations detected contained TE sequences but none had boundaries that matched TE boundaries (“TE = SV”), suggesting that the polymorphisms were the result of deletions, not TE insertions (Figure 4).

**Figure 3.**
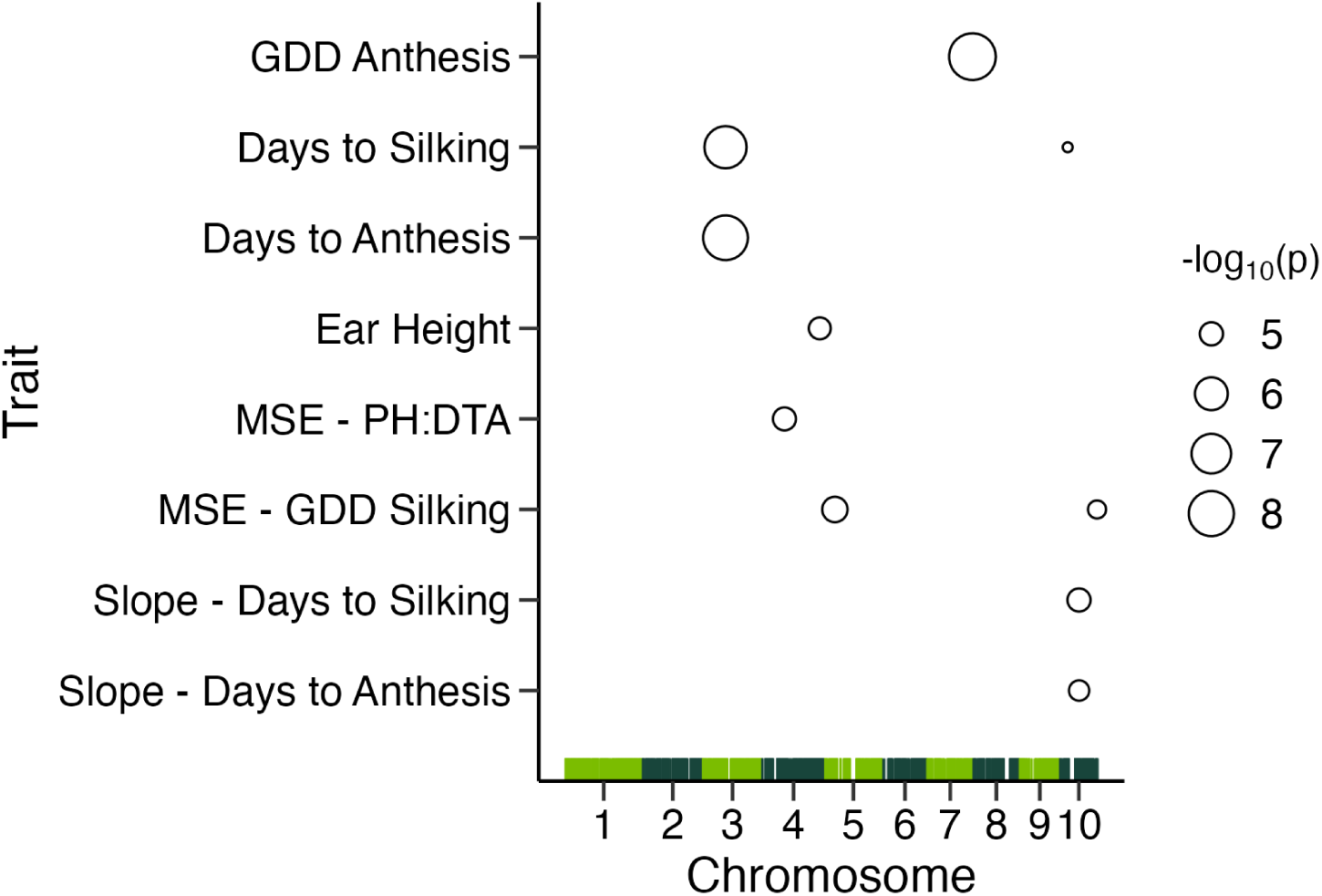
Genomic positions and p-values for eight traits and nine markers with significant SV presence/absence associations. – Bars along bottom represent the genomic positions for the 3,087 SV markers used in the association panel, with chromosomes in alternating colors. Points are sized according to the −*log*_10_*(p)* (GDD: growing degree days; MSE: mean squared error; PH:DTA: ratio of plant height to days to anthesis). Note that the same SV was associated with Days to Silking and Days to Anthesis so there are 10 points total.

**Figure 4.**
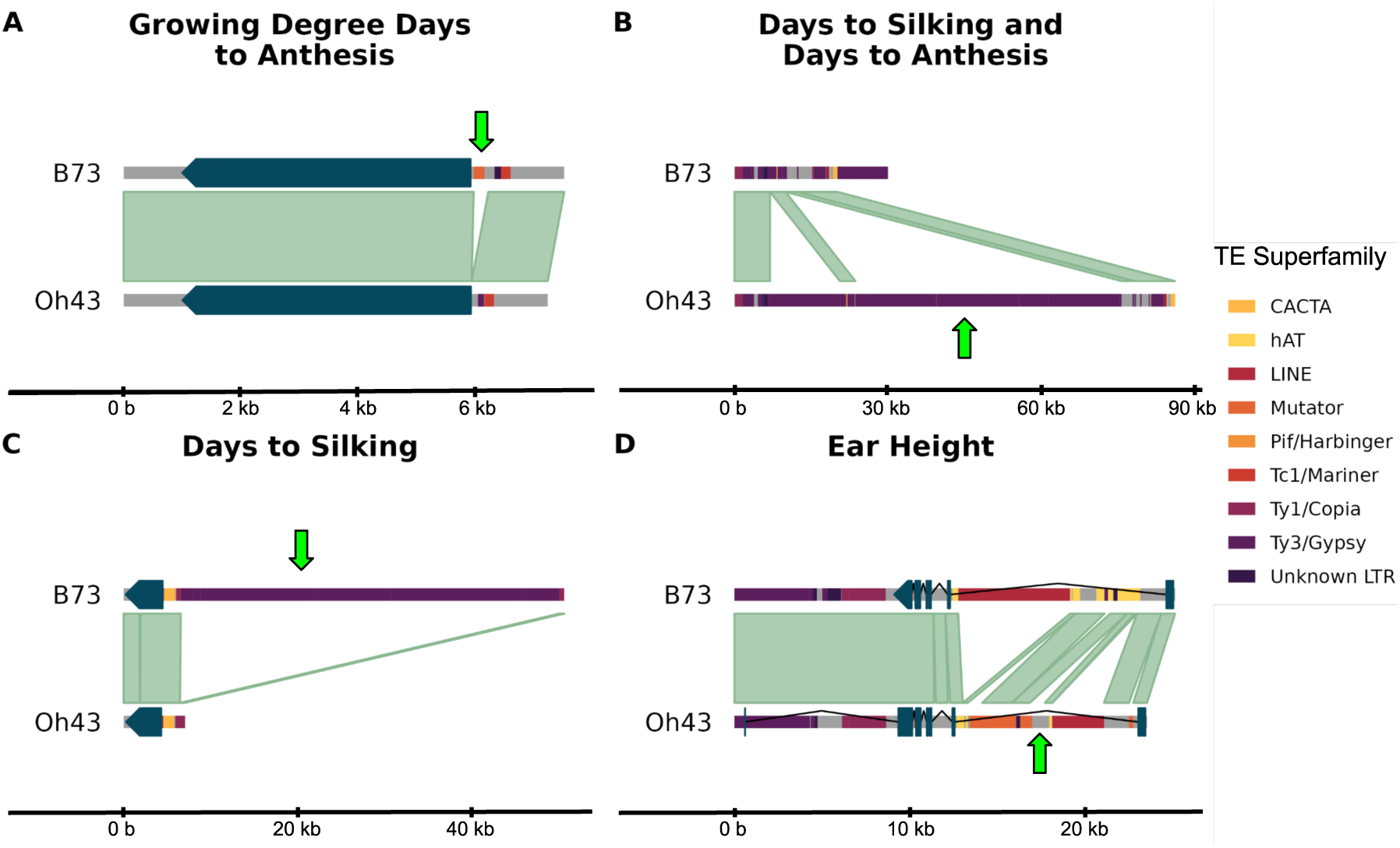
Trait associated structural variant polymorphisms between B73 and Oh43. – Green arrows point to polymorphic SV regions. Alignable regions are shown as green bars between genotypes. TEs are displayed inline and, therefore, do not display overlapping or nested TEs. **(A)** A mutator TE within an SV is present in B73 and absent in Oh43. This SV is 54 bp upstream of the B73 gene Zm00001eb330210, syntenic with Oh43 gene Zm00039ab336990. **(B)** A large SV containing multiple Ty3/Gypsy TEs is present in Oh43 and absent in B73. This intergenic SV is approximately 215 kb from the nearest gene. **(C)** A multi TE SV composed entirely of Ty3/Gypsy TEs is present in B73 and 2091 bp upstream of the gene Zm00001eb411130 (syntenic with Oh43 gene Zm00039ab420040). **(D)** A polymorphic incomplete TE - SV is located within the Oh43 gene Zm00039ab208360 is present in Oh43 and absent in B73.

The SV associated with growing degree days to anthesis is within B73 on chromosome seven, 54 bp upstream of the B73 gene Zm00001eb330210 (syntenic with Oh43 gene Zm00039ab336990) (Figures 3, 4A). There are no currently known functions for these genes in maize, nor their orthologs in other species including sorghum, foxtail millet, rice, or *Brachypodium distachyon*. There is evidence of increased expression in these genes in maize in whole seed, endosperm, and embryo for most 2-day increments post pollination (Walley *et al*., 2016). This SV contained a mutator TE within it, but the SV boundaries did not match the TE boundaries.

One SV polymorphism was associated with both days to silking and days to anthesis. This SV is present on chromosome three in Oh43 and is a large, ∼52 kb multi-TE SV composed primarily of Ty3/Gypsy elements (Figures 3, 4B). This region is nearly 215 kb away from the nearest gene. An additional SV associated with days to silking is located on chromosome ten and contains ∼43.5 kb of multiple Ty3/Gypsy TEs (Figures 3, 4C). This SV, present in B73 and absent in Oh43, is 2,091 bp upstream of the gene Zm00001eb411130 (syntenic with the Oh43 gene Zm00039ab420040). Zm00001eb411130, which is also called ZmMM1, is a MADS-box gene and is orthologoues with the OsMADS13 gene in rice and the STK gene in *Arabidopsis thaliana*. OsMADS13’s expression in rice is restricted to the ovule and controls both ovule identity and meristem determinancy during ovule development (Lopez-Dee *et al*., 1999; Dreni *et al*., 2007; Li *et al*., 2011). Similar to OsMADS13, STK in *Arabidopsis thaliana*, which encodes for a MADS-box transcription factor, is expressed in the early floral development in the ovule. Additionally, STK determines ovule identity and also regulates a network of genes that controls seed development and fruit growth (Mizzotti *et al*., 2014; Di Marzo *et al*., 2020). Both OsMADS13 and STK are members of the D-class genes in the ABCDE model for floral development.

The SV associated with ear height contains a partial sequence of a mutator DNA transposon and is on Oh43 chromosome four within an intron of gene Zm00039ab208360 (syntenic with B73 gene Zm00001eb203840) (Figures 3, 4D). This gene, also called *traf42*, is a tumor receptor-associated factor (TRAF) and codes for a BTB/POZ domain-containing protein *POB1*. Although TRAF domain containing proteins are ubiquitous across eukaryotes, there are far more genes encoding TRAF domains in plants compared to animals (Oelmüller *et al*., 2005; Cosson *et al*., 2010). In maize, *traf42* mediates protein-protein interactions (Dong *et al*., 2017) and mutations in the maize gene ZmMAB1, which contains a TRAF domain and is exclusively expressed in the germline cause chromosome segregation defects during meiosis (Juranić *et al*., 2012). Additionally, *POB1* is involved in drought tolerance in the Antarctic moss, *Sanionia uncinata* (Park *et al*., 2018).

### SV genotypes are associated with GxE

We detected five significant associations (FDR *<* 0.05) between SV presence/absence and one of two measures of plasticity (FW regression slope and MSE) for four of the eleven traits tested: the ratio of plant height and days to anthesis (MSE), growing degree days to silking (MSE), days to silking (slope), and days to anthesis (slope)(Figures 3, S9). Four of the five SVs identified contained TE sequence and two SVs appeared to be directly caused by TE insertions.

On chromosome four, we detected an association between an SV and the MSE of the ratio of plant height to days to anthesis across growing locations. This SV appeared to be caused by a partial deletion of a Ty3-like LTR retrotransposon and was not proximal to any gene models in either the Oh43 or B73 alignments.

On chromosome five, we detected an association between an SV and the MSE of growing degree days to silking across growing locations. This SV appeared to be caused by a partial deletion of a hAT TIR transposon but was not proximal to any gene model in either the Oh43 or B73 alignments.

On chromosome ten, we detected three association between SVs and plasticity: the slope of days to silking, the slope of days to anthesis, and the MSE of growing degree days to silking. The SVs associated with days to silking appeared to be the direct result of insertions of hAT TIR transposons, the SV associated with the MSE of growing degree days to silking appeared to an insertion of a PIF Harbinger TIR transposon, but the SV associated with the slope of days to anthesis did not contain TE sequence. The SV associated with the slope of days to silking was 713 bp from the uncharacterized Oh43 gene Zm00039ab424300 (a syntelog of B73 gene Zm00001eb415280), while the SVs associated with the slope of days to anthesis and the MSE of growing degree days to silking were not proximal to any B73 or Oh43 gene model.

### SV genotypes are associated with differential gene expression

We tested for associations between the genotypes of the nine SVs identified by GWAS and gene expression data from three tissues and detected associations for 29 genes (Figure 5). Differentially expressed genes were not immediately proximal to the SV markers they were associated with (the closest differentially expressed gene was 911kb from the associated SV marker) and most were on different chromosomes. Of the 29 significantly associated genes, three genes present in the B73v3 reference alignment were not present in the B73v5 alignment and were removed from further consideration. Of the 26 remaining genes, 11 were associated with a single SV marker on chromosome 10 for the MSE of growing degree days to silking, which was coded as “TE = SV”. The remaining six SV markers identified were associated with between one and four differentially expressed genes and of those six markers, three contained complete TE sequences, two contained incomplete TEs, and one did not contain any TE sequence. Of the three tissues tested, 16 genes were significantly differentially expressed solely in shoot tissue, seven in the the tip of L3, two in the base of L3, and one was differentially expressed in both the shoot tissue and the base of L3.

**Figure 5.**
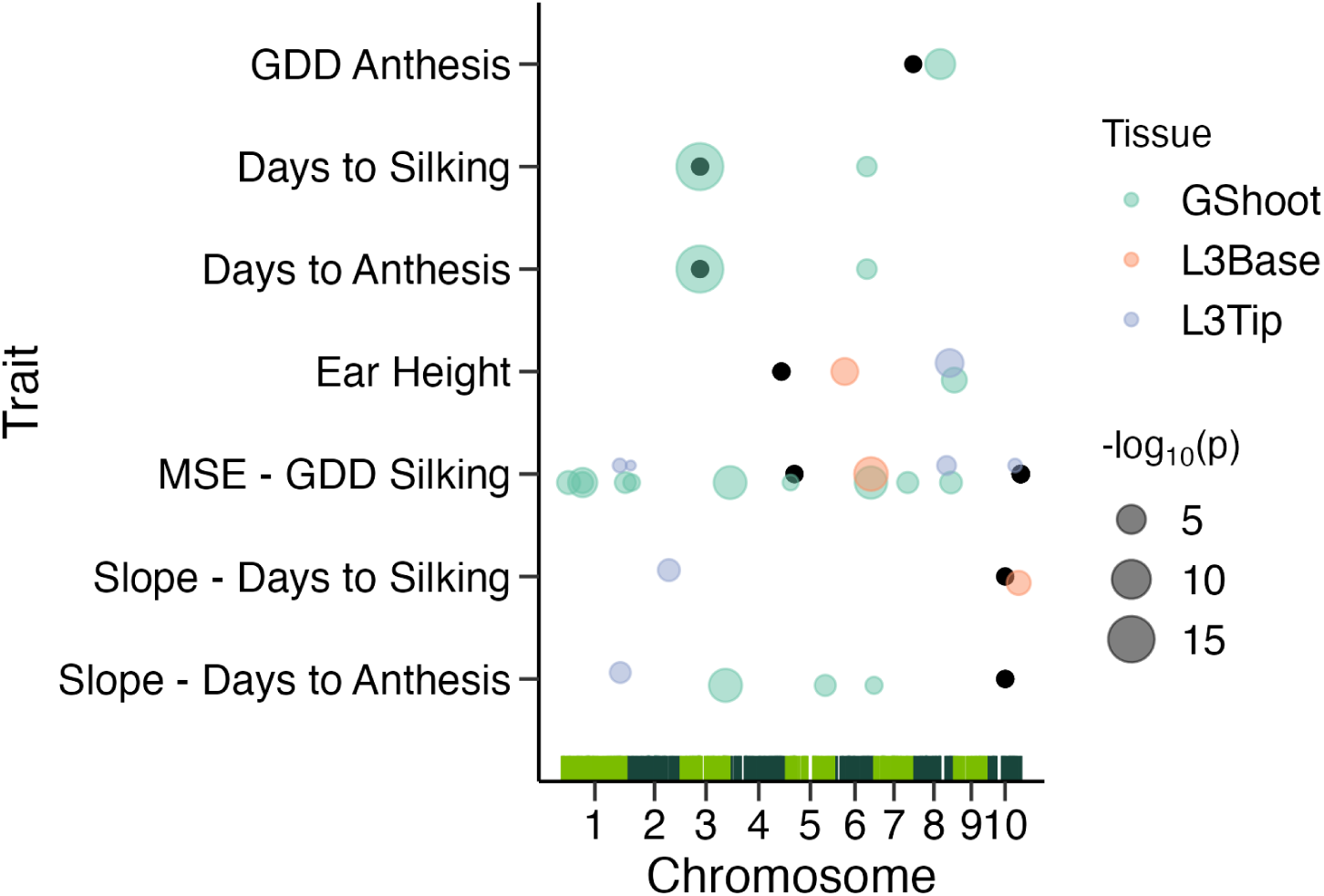
Genomic positions and p-values for genes with expression significantly associated with the genotypes of seven structural variant (SV) markers identified in our genome wide association analyses. – Bars along bottom represent the genomic positions for the 3,087 SV markers used in the association panel, with chromosomes in alternating colors. Black points show the position of the SV marker identified in each trait. Colored points are sized according to the false discovery rate adjusted −*log*_10_*(p)* with tissue collected from germinating shoot (GShoot) in green, the base of leaf three (L3Base) in orange, and the tip of leaf three (L3Tip) in blue. The SV marker on chromosome three was the most proximal to the identified SV marker, but was still 911 kb away (GDD: growing degree days; MSE: mean squared error).

### Most SVs are in linkage disequilibrium with SNPs

All SV alleles used in the GWAS are within 1 Mb (mean distance of 649 bps) from the nearest SNP present in the HapMap3 dataset (Figure S10) and all SVs have an *r*^2^ *>* 0.1 with at least one nearby SNP. Only 6 SVs had an *r*^2^ *<* 0.5 with any nearby SNP. For the SV alleles that are significant to traits, all have a SNP in perfect LD.

Despite high LD between SVs and nearby SNPs, many of the associations detected between SVs and traits would not have been captured with a GWAS using all SNPs. Of the four SVs associated with main effects, only one was found in the same peak regions in the SNP GWAS (Figures S11, S12). This lack of overlap between the SV GWAS associations and the SNP GWAS associations is a result of different significance cutoffs in the two different analyses. The HapMap3 SNP dataset used in the GWAS has 16,435,136 SNPs while there were only 3,087 SVs in the SV association mapping analysis, so a SNP needed to have a p-value below 7.94 × 10^−6^ (averaged across traits) to overcome the FDR cut-off in the SNP GWAS while its linked SV only needed a p-value below 1.86 × 10^−4^ (averaged across traits) to be detected as significant in the SV GWAS.

## Discussion

In this study we leveraged two reference genomes along with a broader set of short-read genomic data to capture SV diversity in a maize diversity panel. The maize genome’s highly repetitive nature makes it challenging to rely on short-read alignments alone to characterize SV polymorphism *de novo* (Hufford *et al*., 2021). By ascertaining SVs presences and absences between two genotypes, we were able to call SVs across hundreds of maize genotypes using short-read data and identify SVs associated with trait variation.

We found nine SV polymorphisms associated with either average trait value or trait plasticity in a variety of maize phenotypes (Figure 3). Previous studies have identified SVs associated with phenotypic variation that would not be discovered in analyses that use SNPs alone (Yang *et al*., 2019; Guo *et al*., 2020; Hartmann, 2022; Zhang *et al*., 2024). Here, while the SV GWAS identified hits that were not present in the SNP GWAS, all SV associations detected were in perfect linkage disequilibrium with SNPs. We did not detect associations that were not captured by the SNP dataset but instead these SVs reached statistical significance because there were many fewer SVs than SNPs. Previous work investigating TE polymorphism in a different maize genetic diversity panel did find that 20% of TEs were not in LD with SNPs but these SNPs tended to be at a low minor allele frequency in the population (Qiu *et al*., 2021b). By focusing on common SV polymorphisms we likely have missed many SVs that are low frequency and not in LD with surrounding SNPs – however these low frequency SVs would be unlikely to be associated with trait variation in a GWAS.

Of the SVs included in this study, 91% contained TEs or are themselves of TE origin and the largest category of SVs were clear examples of TE insertion (21,103 or 23.5%). All but one of the SVs associated with trait variation and with GxE contained TE sequence, yet only the SVs on chromosome ten for the slope of days to silking and the MSE of growing degree days to silking FW models appeared to be the direct result of TE insertions. The remaining seven associations result from deletions that contain TEs. This result is consistent with previous findings that deletions have been the dominant contributors to SV polymorphism in maize (Munasinghe *et al*., 2023). We did observe that the SV associated with the MSE of growing degree days to silking on chromosome ten that appeared to result from a TE insertion was the SV with the most associations with gene expression. This pattern is consistent with hypotheses that TE insertions are particularly likely to affect gene expression (Klein and Anderson, 2022), although further work is clearly needed to evaluate how broad this pattern is across a larger sample of SVs.

We found five significant associations between SVs and plasticity, quantified using mean squared error and slopes from the Finlay-Wilkinson regression models. The finding that different SVs were associated with traits than with trait plasticity is consistent with most previous work. For example, the genetic architecture of trait means and trait plasticity have been shown to differ in maize (Kusmec *et al*., 2017; Tibbs-Cortes *et al*., 2024) and *Arabidopsis thaliana* (Fournier-Level *et al*., 2022) but not sorghum (Wei *et al*., 2024). We also did not see a clear pattern that SVs are more likely to affect trait variation across environments than trait means, but this may result from having a small number of associations across both categories.

Overall, we have demonstrated an approach for using two reference genomes to identify structural variants and then genotype for these variants in a larger panel of individuals with short-read sequencing data. This approach identifies SVs associated with phenotypic variation and with GxE interactions. However, this approach does bias us towards common alleles that were polymorphic within the two reference assemblies. This bias is acceptable for a GWAS, where we will also be biased towards detecting associations with variants at intermediate allele frequency, but would be less appropriate for any analysis that would need to identify SVs with low allele frequencies. As long-read data becomes more affordable and more reference genomes become available for more species, these types of approaches will improve our ability to detect SVs and investigate their potential functional importance.

## Data availability

All sequencing data used are publicly available and generated by previous papers.

## Acknowledgments

We thank Nathan Springer, Jeff Ross-Ibarra, Michelle Stitzer, and Yaniv Brandvain, along with members of the Josephs, Hirsch, Ross-Ibarra, Kaepler, and Springer labs for helpful comments and suggestions.

## Data Accessibility Statement

All data used in this paper came from publicly available databases. We have included information for accessing data resources in the appropriate places of the Materials and Methods section. A github repository with all code and a table of TE polymorphisms will be made available upon publication and archived at zenodo.

### Benefit-sharing Statement

All code and a table of called TE polymorphisms will be made available, as described above

## 1 Supplementary Information

**Figure S1.**
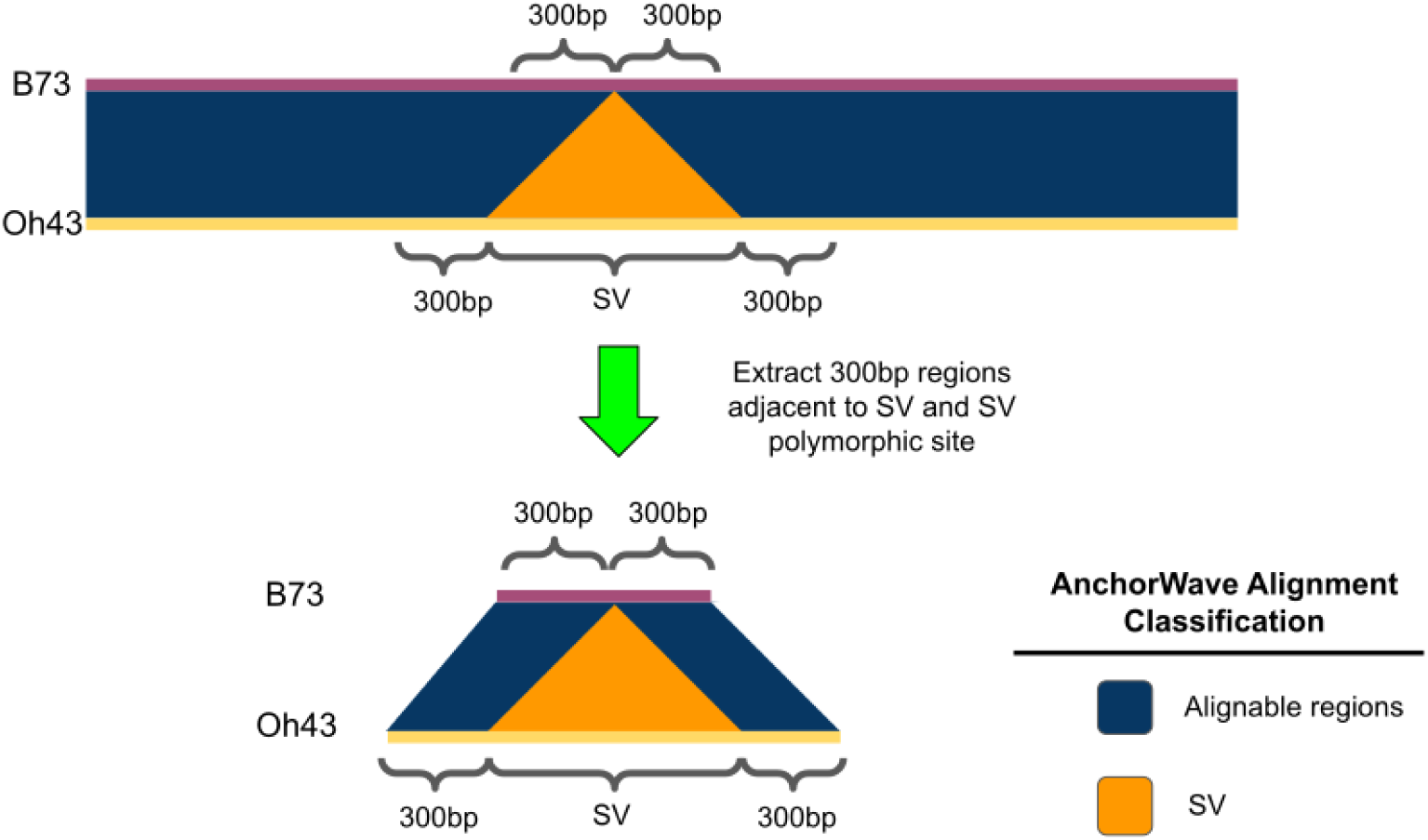
Extracting SV present and absent alleles. – For each polymorphic SV between B73 and Oh43 identified in Munasinghe, et al. (2023), we extracted 300 bp flanking alignable regions along with the SV for to make “SV present alleles” while 300 bp flanking alignable regions were extracted around the insertion point, which we term “SV-absent alleles”.

**Figure S2.**
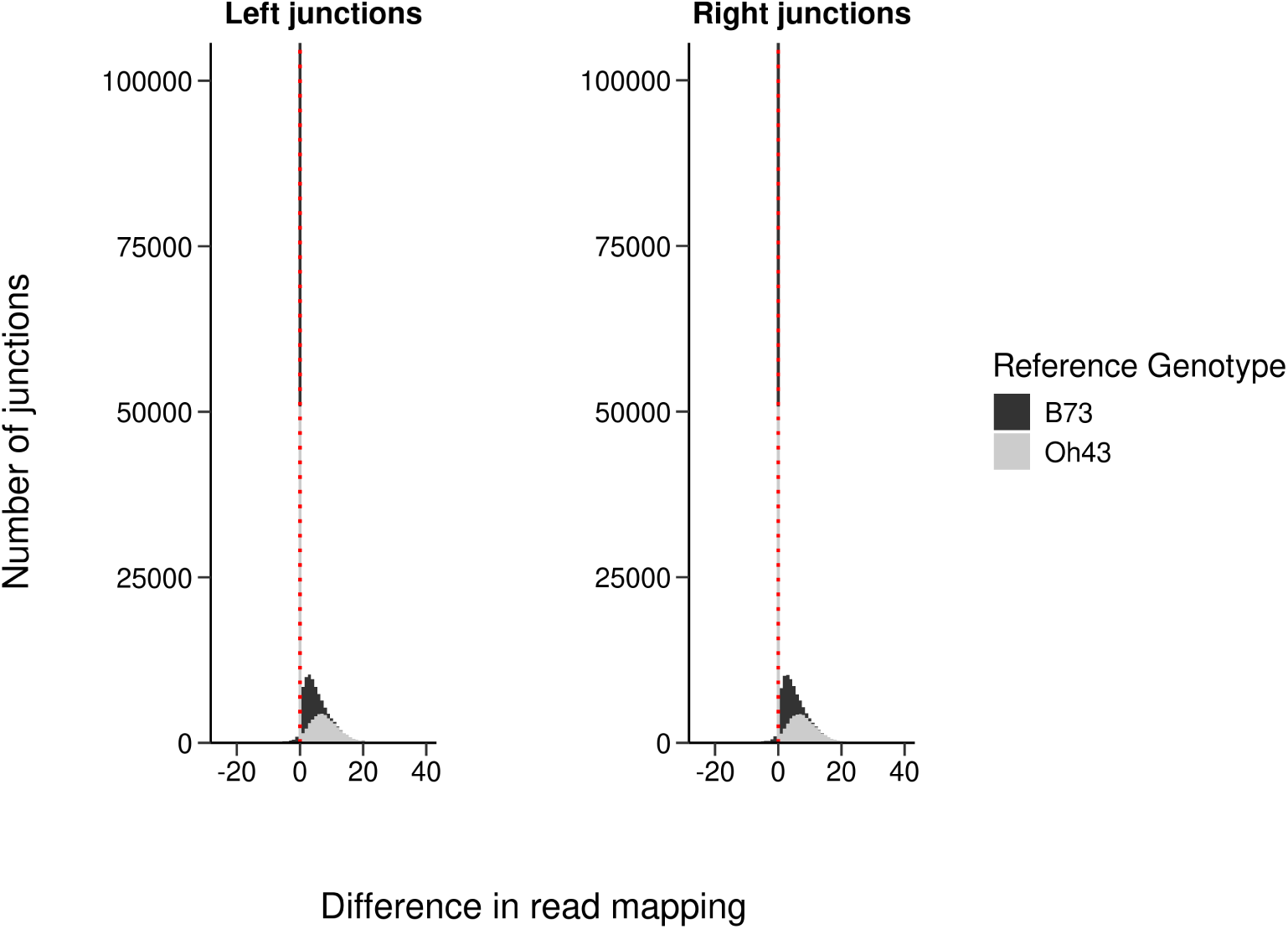
Read mapping differences at the left and right junctions for all SVs. – Differences were calculated as the number of reads from the non-parent genotype subtracted from reads mapping from the parent genotype. A positive difference indicates SVs that are supported and retained for future analyses.

**Figure S3.**
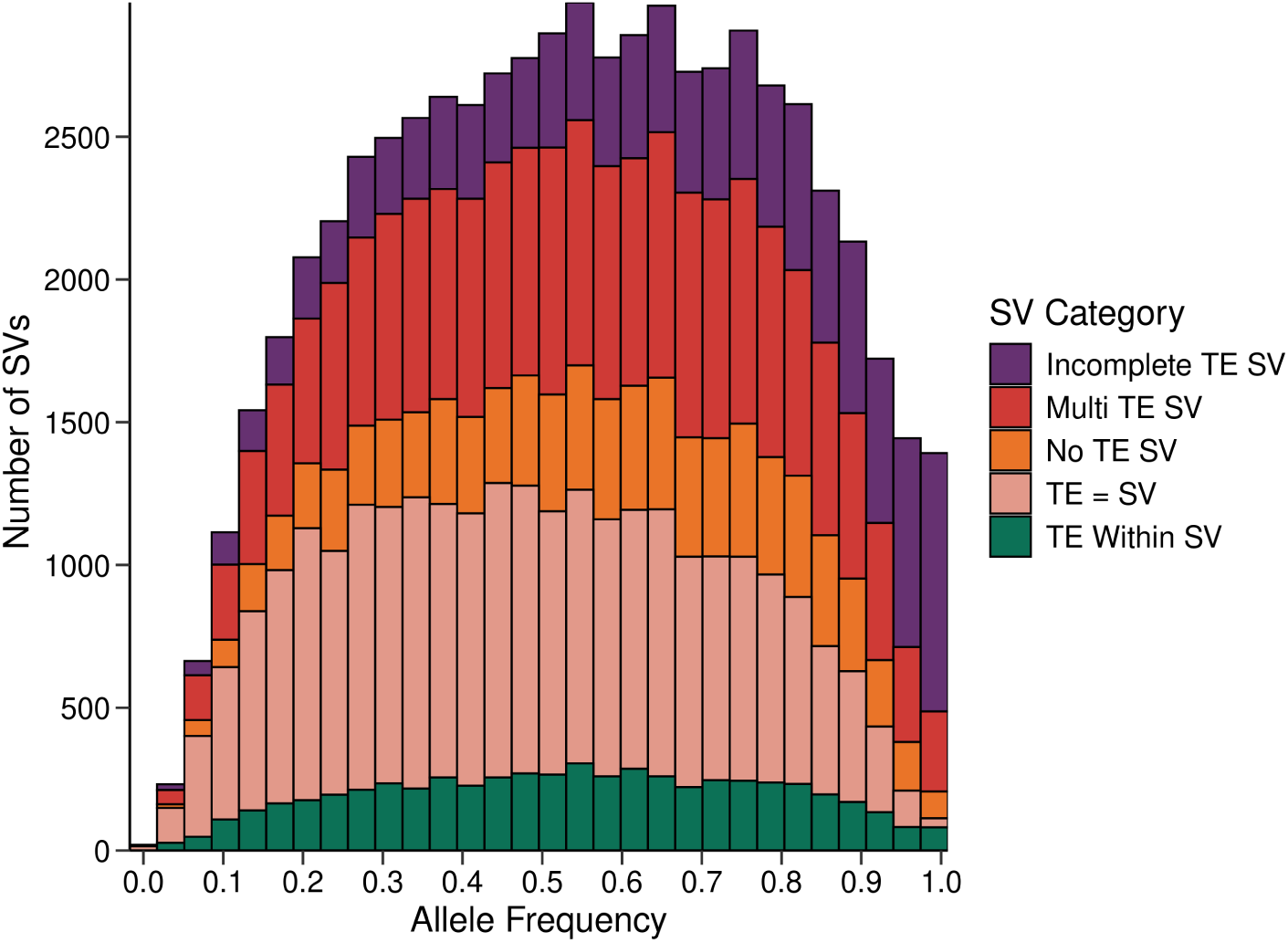
SFS of SVs categorized by TE content. – “Incomplete TE SV” and “Multi TE SV” categories SV polymorphisms skew towards moderate to high frequencies whereas all other categories skew towards low to moderate frequencies.

**Figure S4.**
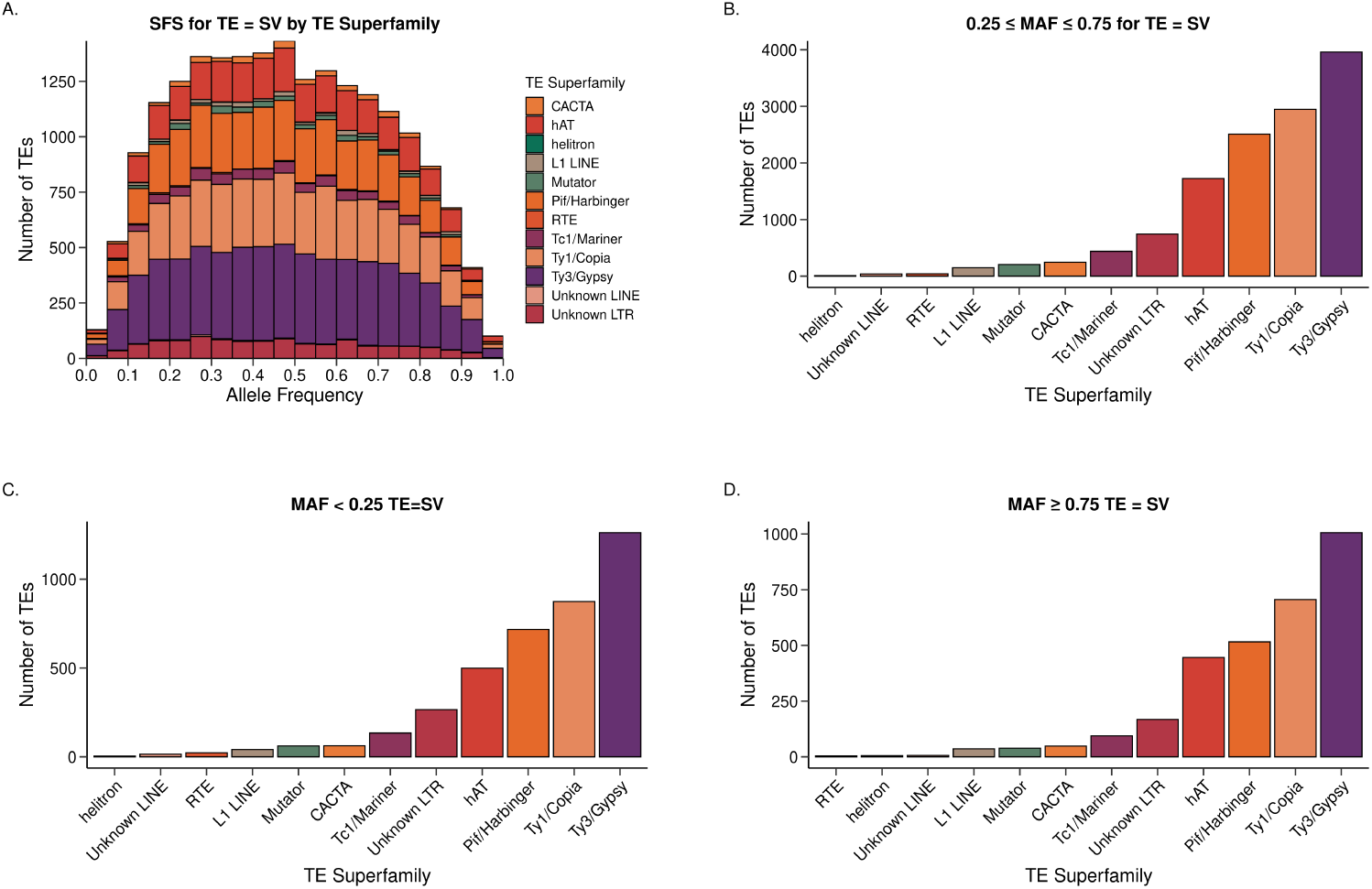
SFS and MAF of TE superfamilies for TE = SV. – (**A**). TE polymophisms skew towards moderate frequency. (**B**), (**C**), and (**D**). Frequencies for all TE superfamilies are consistent across all MAF thresholds.

**Figure S5.**
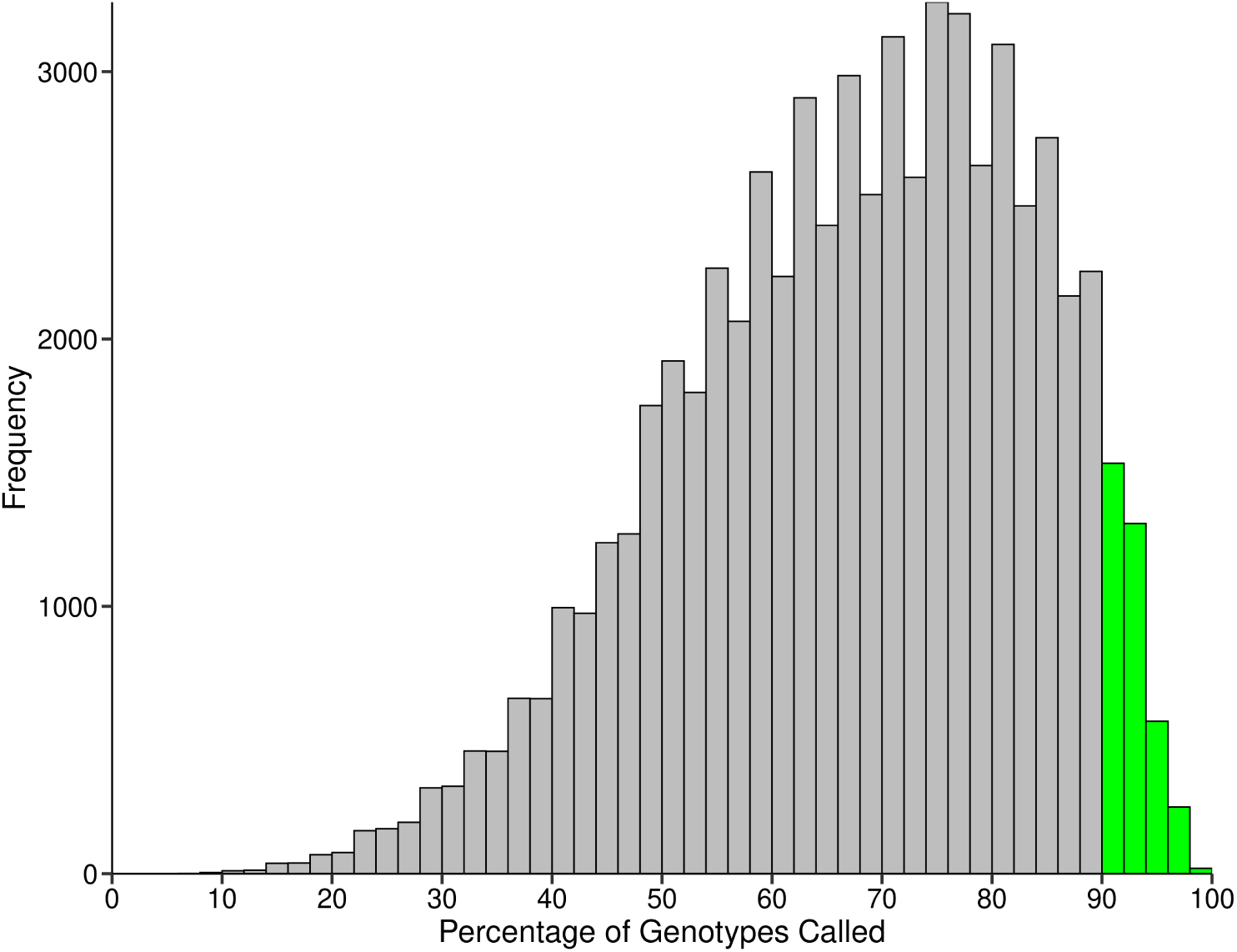
Percentage of genotypes called per SV. – Green bars indicate SVs with at least 90% of genotypes called and are retained for GWAS.

**Figure S6.**
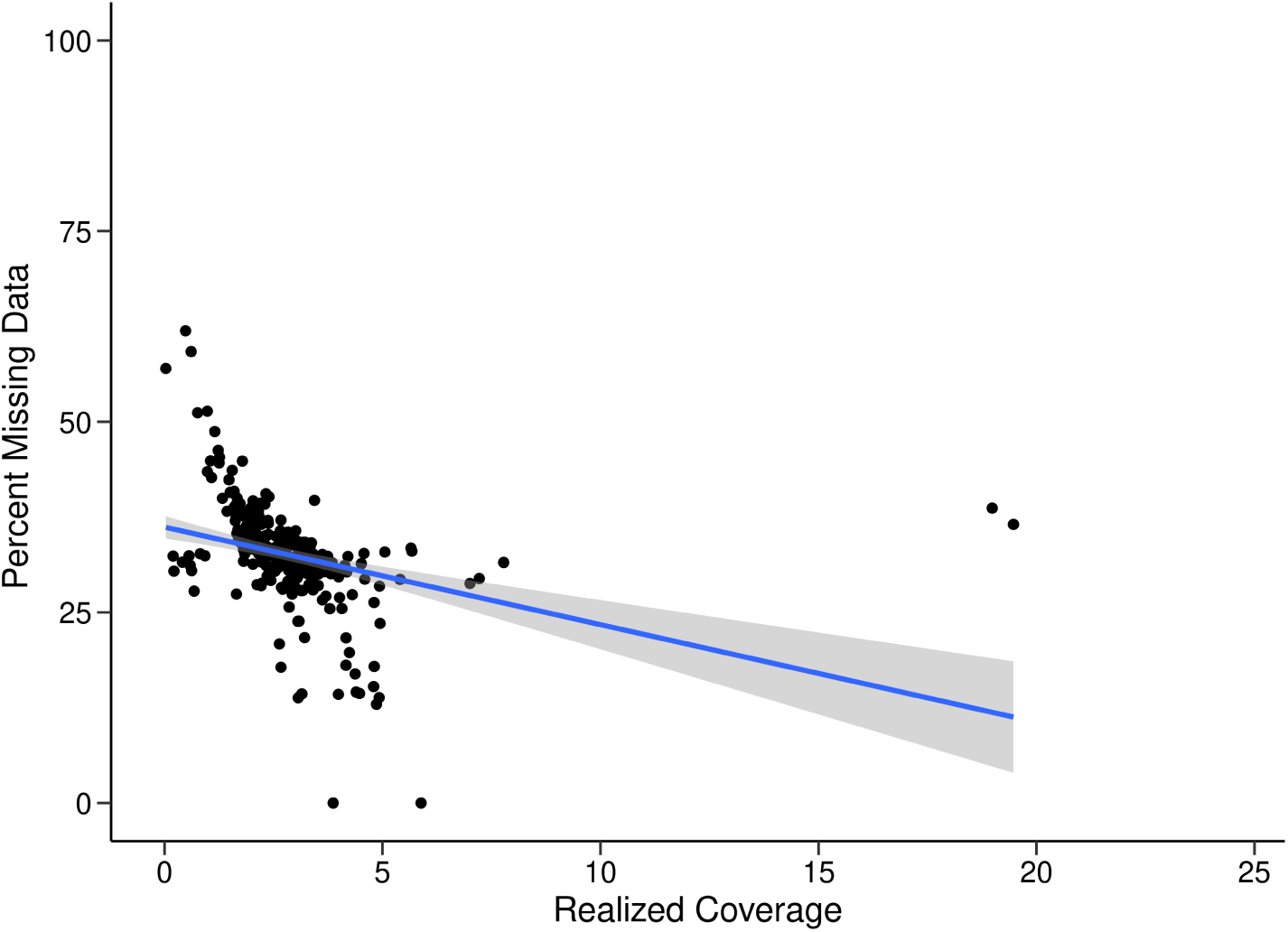
Linear Model of relationship between missing data and read coverage - all genotypes. – Adjusted R-squared: 0.1045, F-statistic: 33.21, p-value: 2.22 × 10^−8^

**Figure S7.**
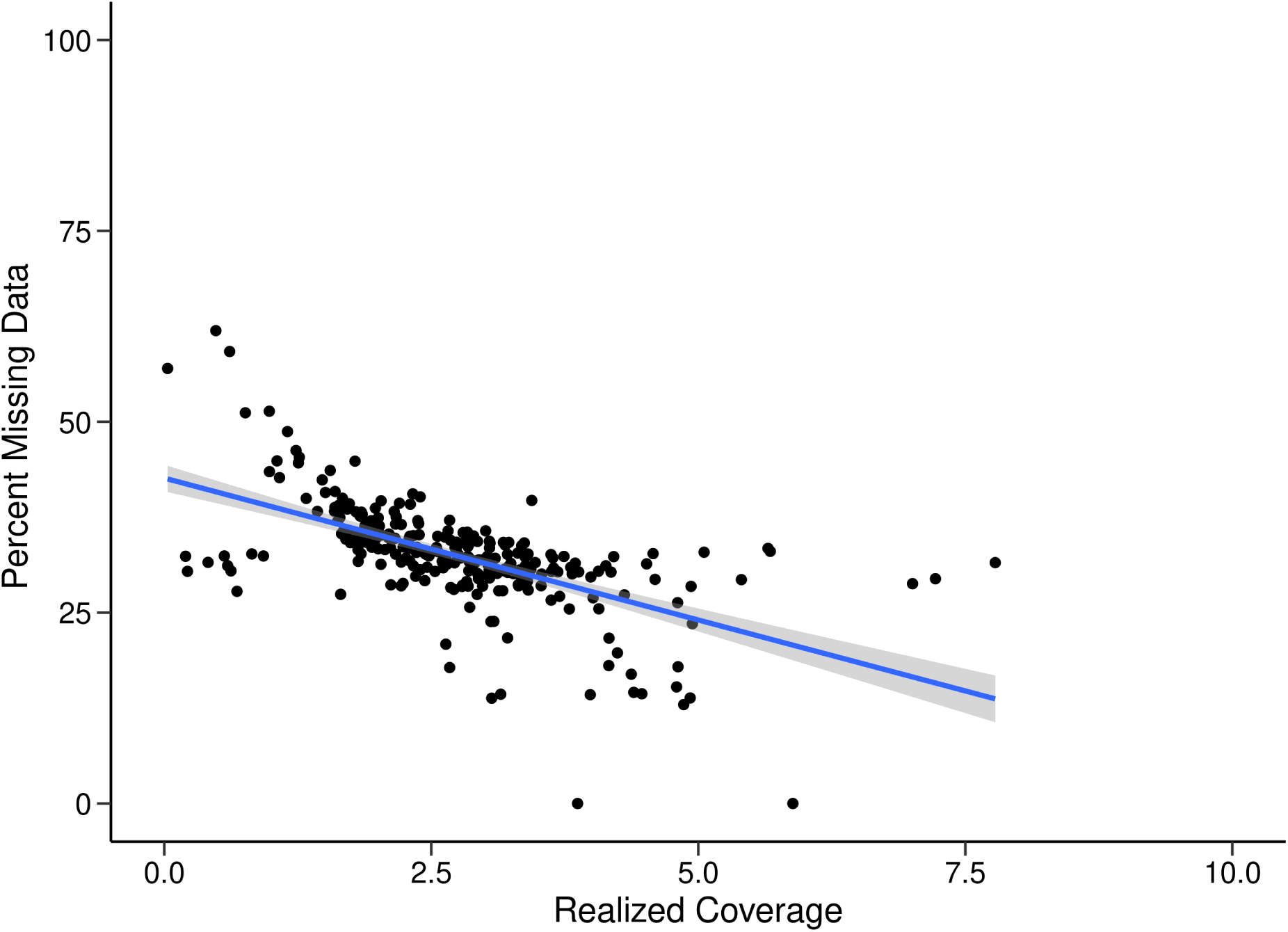
Linear Model of relationship between missing data and read coverage - 2 outliers removed. – Adjusted R-squared: 0.3576, F-statistic: 153.50, p-value: 2.2 × 10^−16^

**Figure S8.**
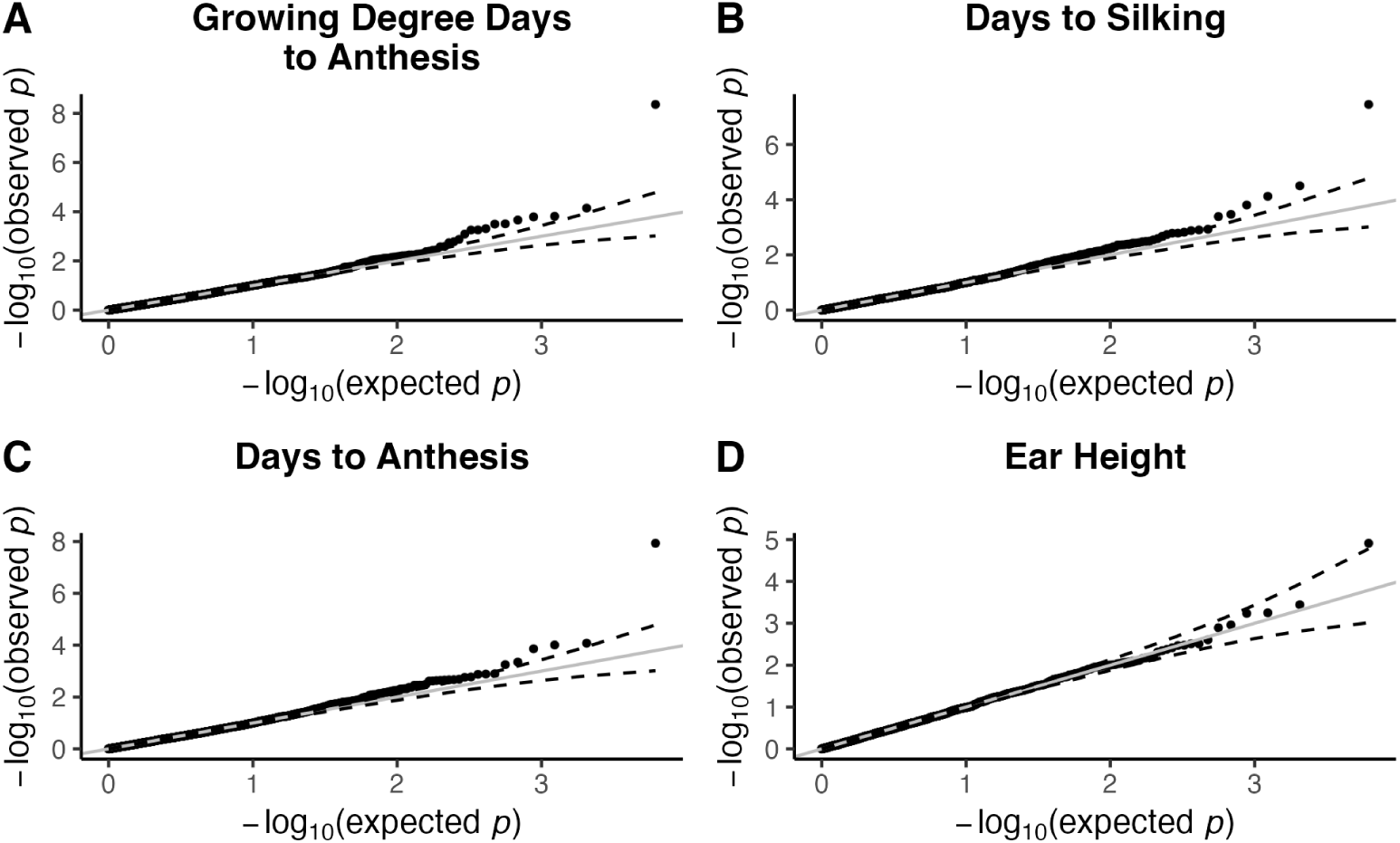
Q-Q plots for traits with SV associations

**Figure S9.**
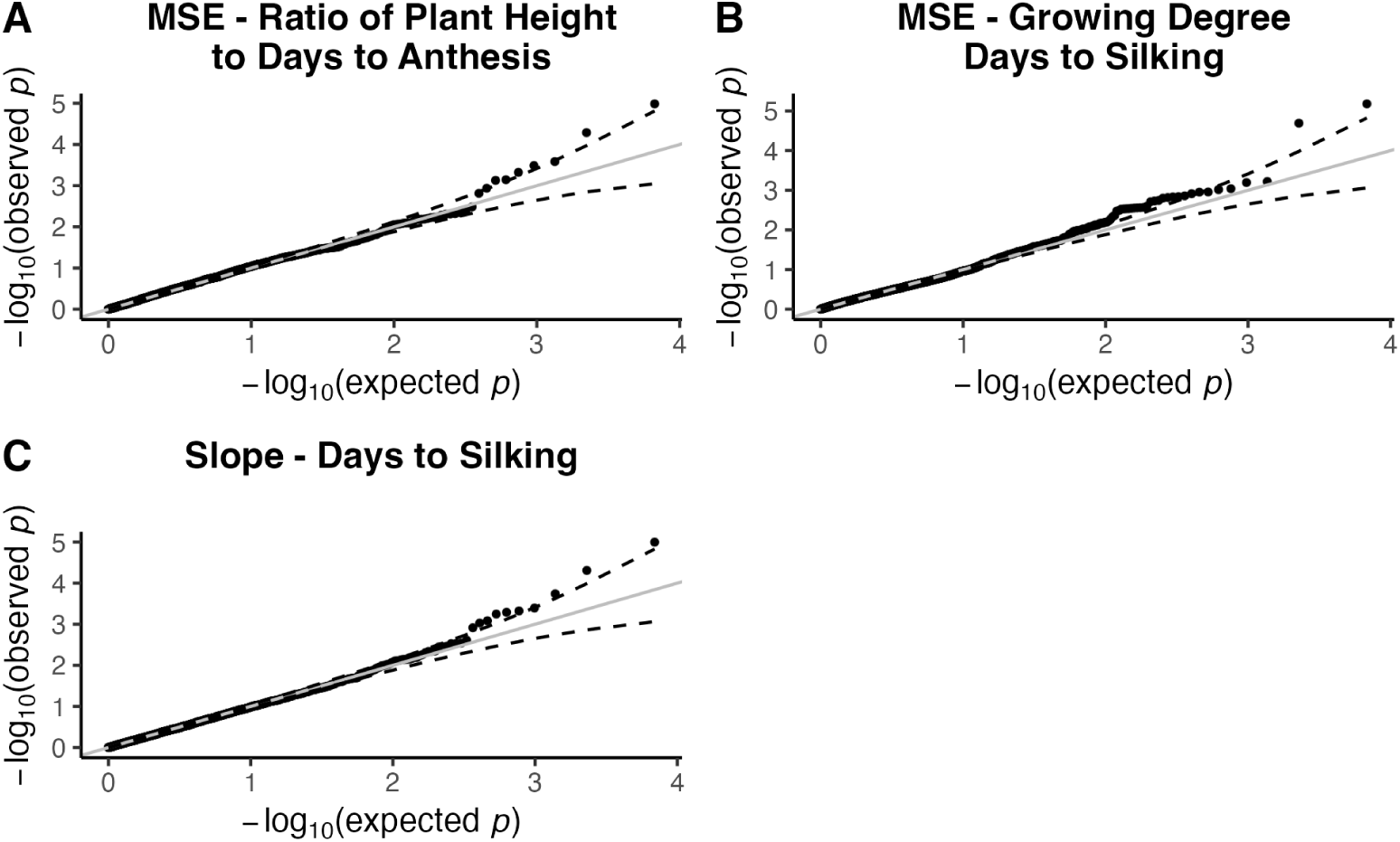
Q-Q plots for Finlay-Wilkinson regression traits with SV associations. – (A). the mean-squared error (MSE) of the ratio of plant height to days to anthesis, (B). the MSE of growing degree days to silking, (C). the slope days to silking. Note the deviations between expected and observed p-values in the MSE of growing degree days to silking model (B).

**Figure S10.**
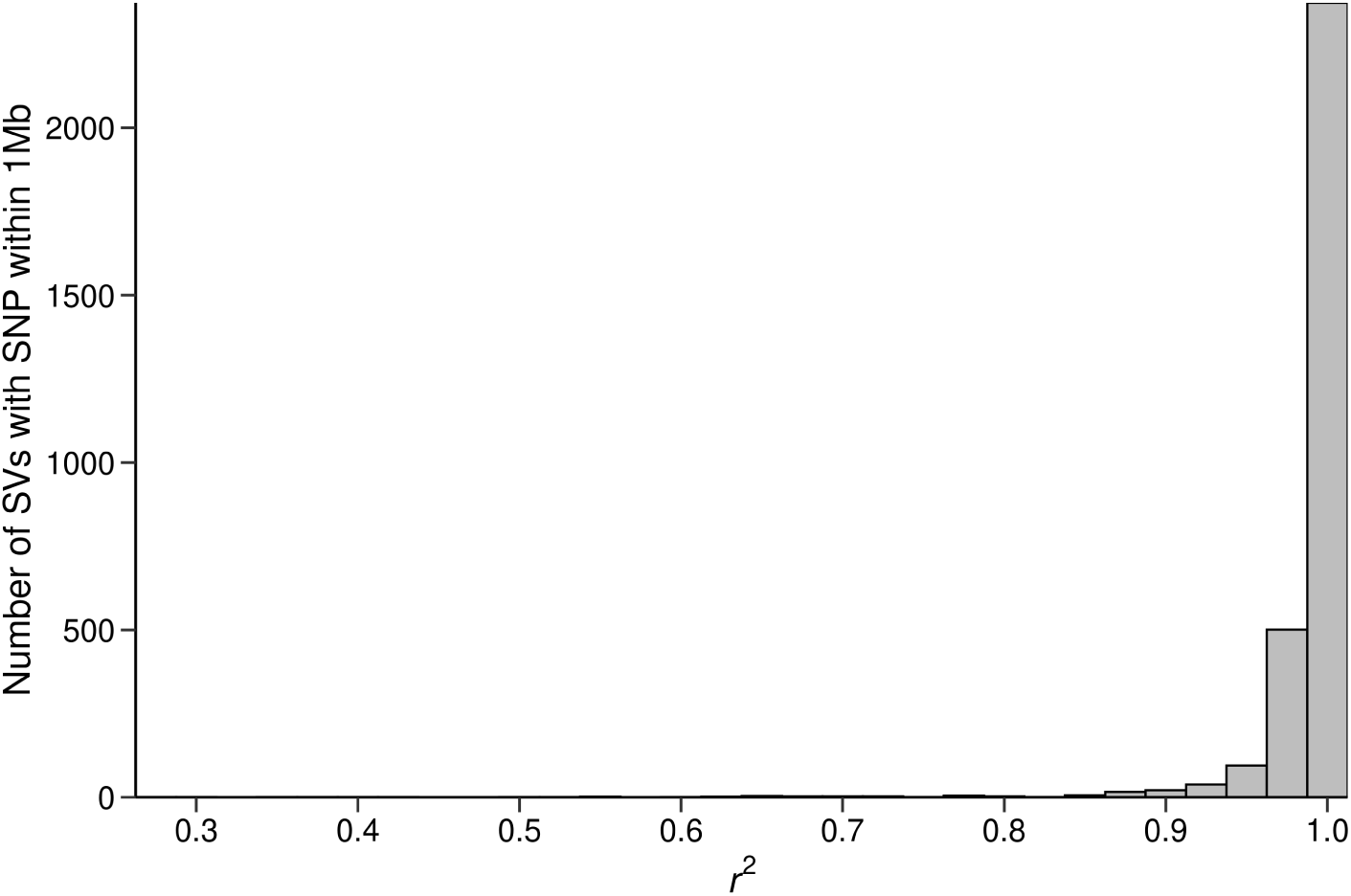
Highest LD between SVs and SNPs within 1Mb. – A large proportion of the 3,087 SVs used in GWAS are linked with adjacent SNPs. SVs with *r*^2^ = 1 : 2, 277, *r*^2^ ≥ 0.5 : 3, 080.

**Figure S11.**
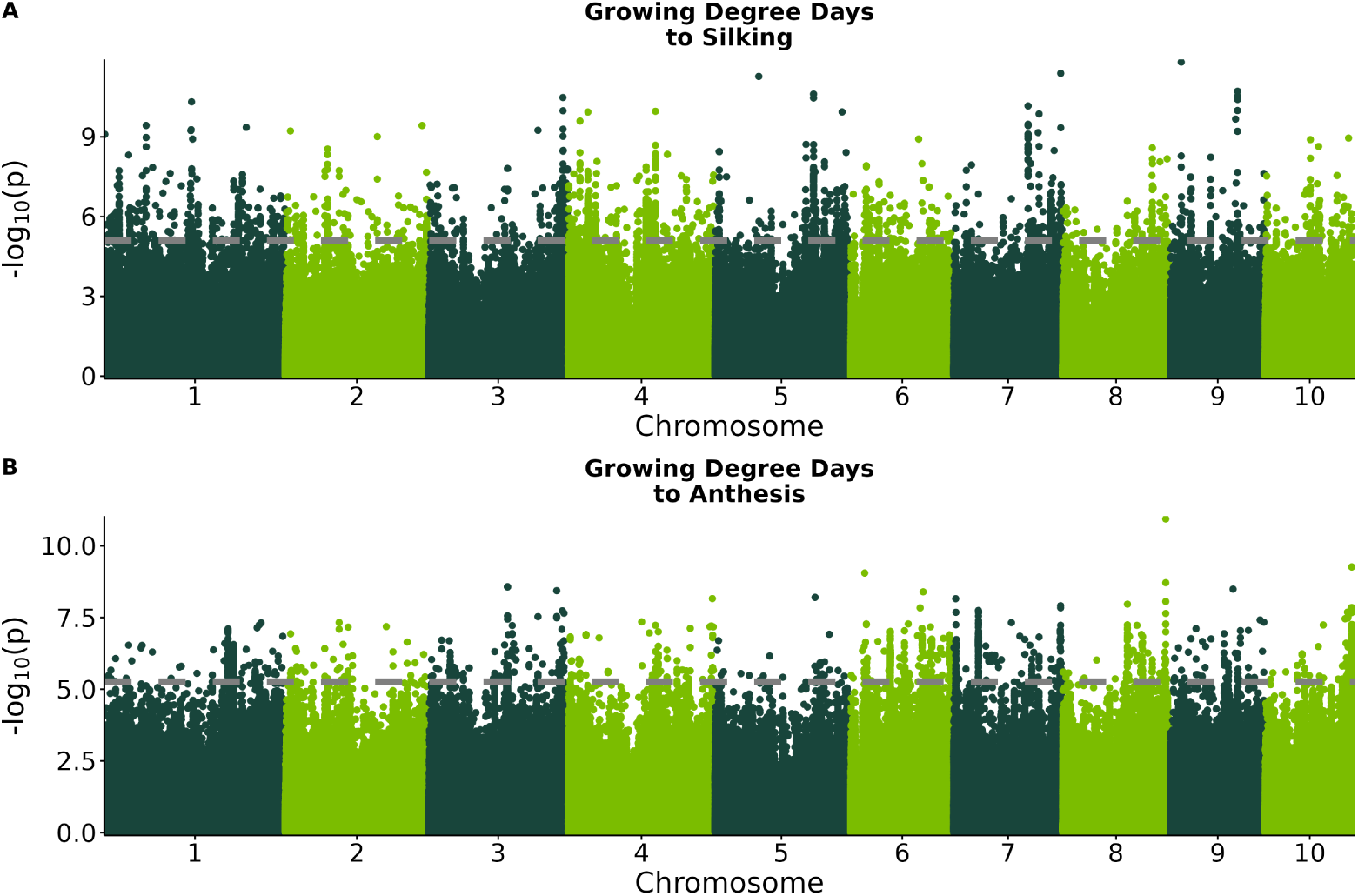
Manhattan Plots of HapMap3 SNPs. – The gray dashed line represents the FDR signifcance threshold. (A) There are several SNPs associated with growing degree days to silking, although none are in LD with SVs associated with the same trait. (B) There are many SNPs throughout the genome associated with growing degree days to anthesis.

**Figure S12.**
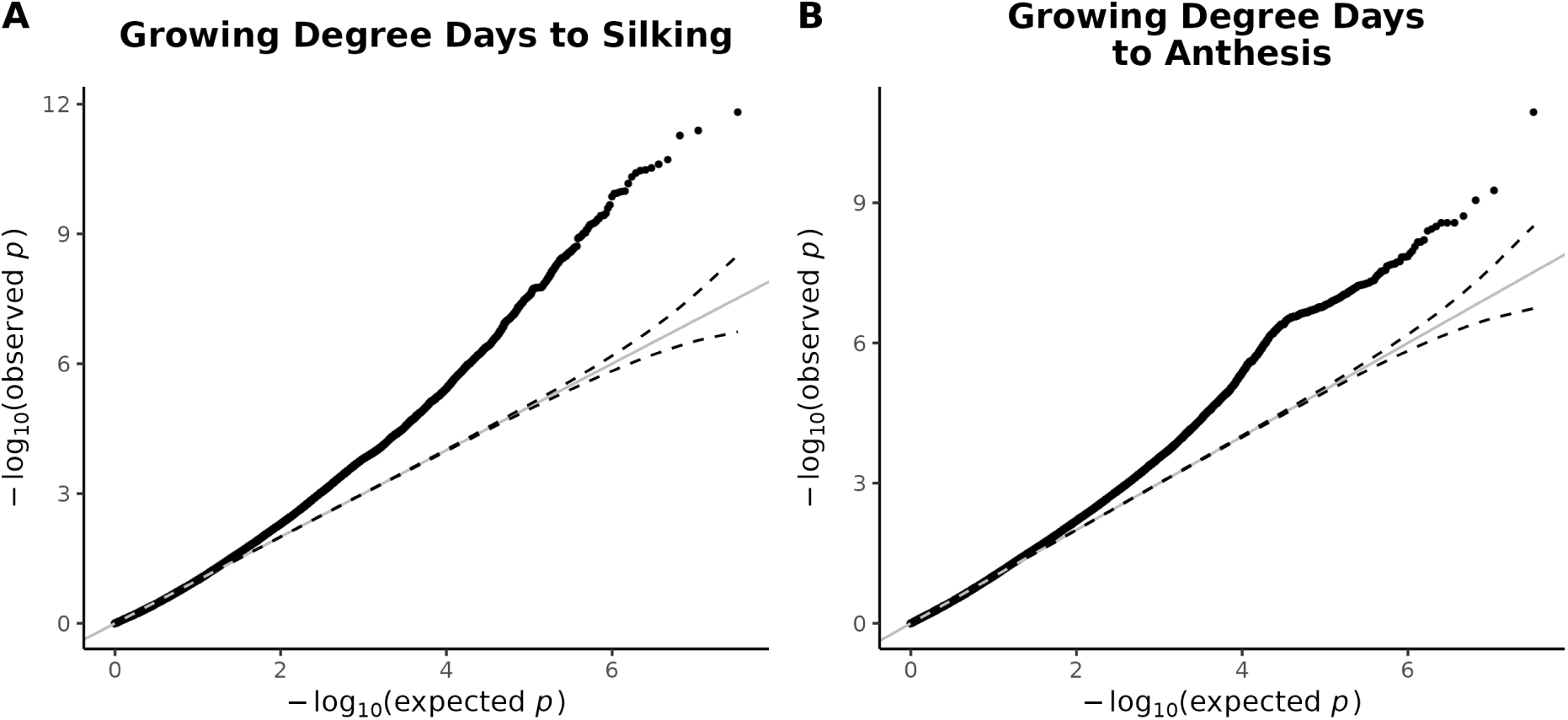
Q-Q plots for traits with HapMap3 SNP associations

